# The edible seaweed *Laminaria japonica* contains cholesterol analogues that inhibit Lipid Peroxidation and Cyclooxygenase Enzymes

**DOI:** 10.1101/2021.10.11.463984

**Authors:** Xingyu Lu, Amila A. Dissanayake, Chuqiao Xiao, Jie Gao, Mouming Zhao, Muraleedharan G. Nair

## Abstract

In this study, 5 sterols were isolated and purified from *Laminaria japonica*, commonly known as edible brown seaweed, and their structures were identified based on detailed chemical methods and spectroscopic analyses. Spectroscopic analyses characterized 5 sterols as 29-Hydroperoxy-stigmasta-5,24(28)-dien-3β-ol, saringosterol (24-vinyl-cholest-5-ene-3β,24-diol), 24-methylenecholesterol, fucosterol (stigmasta-5,24-diene-3β-ol), and 24-Hydroperoxy-24-vinyl-cholesterol. The bioactivities of these sterols were tested using lipid peroxidation (LPO) and cyclooxygenase (COX-1 and −2) enzyme inhibitory assays. Fucosterol exhibited the highest COX-1 and −2 enzyme inhibitory activities at 59 and 47%, respectively. Saringosterol, 24-methylenecholesterol and fucosterol showed higher LPO inhibitory activity at >50% than the other compounds. In addition, the results of molecular docking revealed that the 5 sterols were located in different pocket of COX-1 and −2 and fucosterol with tetracyclic skeletons and olefin methine achieved the highest binding energy (−7.85 and −9.02 kcal/mol) through hydrophobic interactions and hydrogen bond. Our results confirm the presence of 5 sterols in *L. japonica* and its significant anti-inflammatory and antioxidant activity.

**Graphical abstract:** 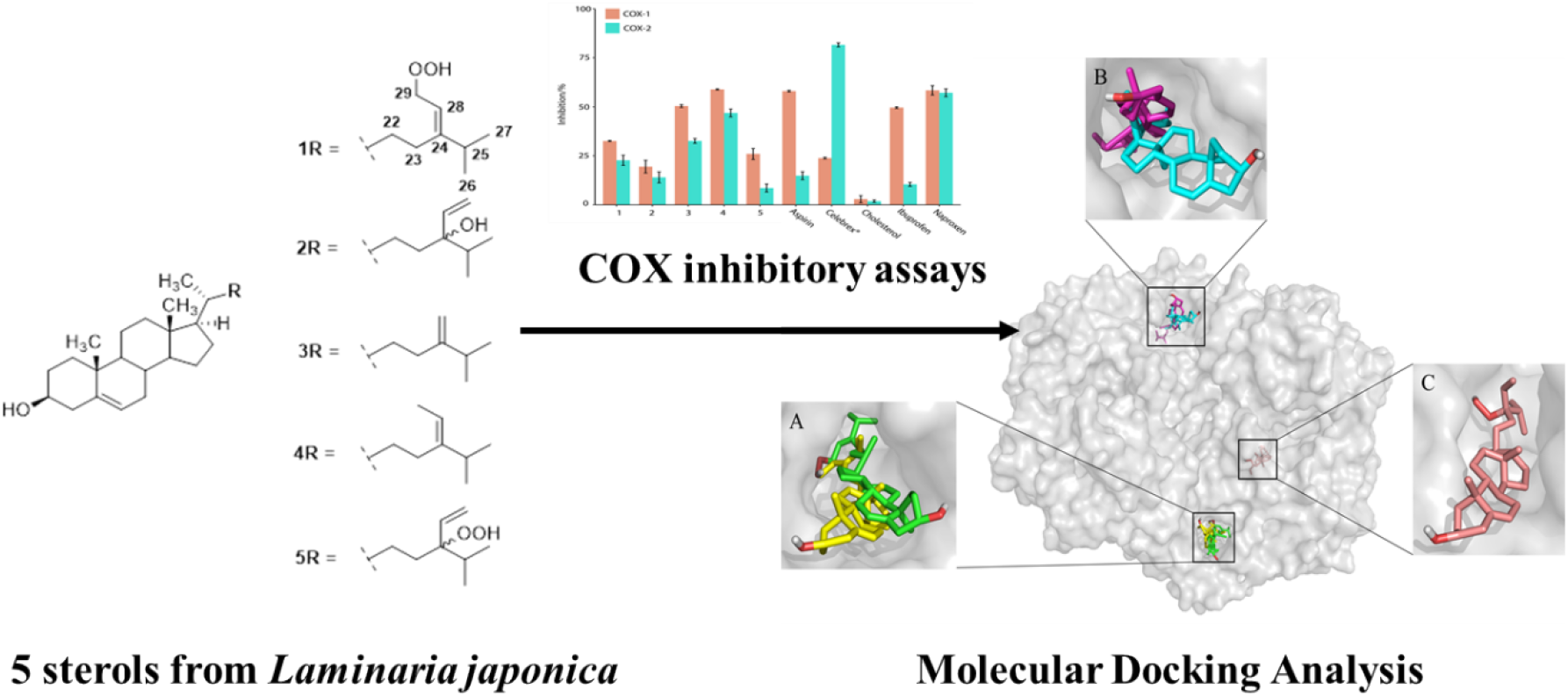

**Highlights:**

1. Sterols 29-hydroperoxy-stigmasta-5,24(28)-dien-3β-ol and 24-hydroperoxy-24-vinyl-cholesterol are identified for the first time in *L. japonica*.
2. Saringosterol, 24-methylenecholesterol and fucosterol showed strong LPO inhibitory activity.
3. Fucosterol showed highest binding affinity for COX-1 and −2 enzymes through hydrophobic interactions.

## 1. Introduction

*Laminaria japonica*, a commercial edible brown seaweed, is widely cultivated and consumed as a health food resource in China, Japan, and Korea. Over the past decades, *L. japonica* is one of the most important marine medicinal foodstuffs and a rich source of various functional compounds, which have been widely investigated by both *in vitro* and *in vivo* experiments.[1–4] Numerous phytochemical investigations on *L. japonica* revealed the presence of polyphenols, fatty acids, carotenoids, steroidal ketones, amine, sterols and tocopherol.[5–13] Fucoxanthin, one of the most abundant carotenoids and the major functional pigment present in *L. japonica*, has been reported to have antioxidative, cytoprotective, anti-inflammatory and anti-obese properties.[6–9] Butyl-isobutyl-phthalate, extracted from *L. japonica*, exhibited hypoglycemic effect in vivo and non-competitive inhibition of α-glucosidase in vitro.[10, 11] Six isolates from EtOAc extracts of *L. japonica* have been determined as 2,6-dibromo-4-(2-(methylamino)ethyl)phenol, 6-bromo-1H-indole-3-carbaldehyde, 1H-indole-3-carbaldehyde, 4-bromobenzoic aldehyde, 4-bromobenzoic acid and 4-hydroxybenzoic acid.[12] Four steroidal ketones including ergosta-4,24(28)-diene-3-one, ergosta-4,24(28)-diene-3,6-dione, stigmasta-4,24(28)-diene-3-one and stigmasta-4,24(28)-diene-3,6-dione have also been isolated from *L. japonica*.[13]

Oxidative stress can activate a variety of transcription factors, which lead to the differential expression of some genes involved in inflammatory pathways.[14] The oxidative stress machinery and inflammatory signaling are not only interrelated, but their impairment can lead to many chronic diseases. Previous phytochemical investigations revealed the hexane extract from *L. japonica* exerts anti-inflammatory effects.[15] Nevertheless, limited studies on the purification of hexane extracts from *L. japonica* and structure characterization of the pure compounds have been performed and, to the best of our knowledge, anti-inflammatory and antioxidant activities of these pure compounds have not been reported. Herein, 5 sterols from hexane extracts of *L. japonica* have been purified by VLC (vacuum liquid chromatography), multi-step MPLC (medium-pressure liquid chromatography), Prep-TLC (preparative thin-layer chromatography) and HPLC (high pressure liquid chromatography), and characterized by high-resolution electrospray ionization mass spectrometry (HRESITOFMS), nuclear magnetic resonance (NMR) and gas chromatography-mass spectrometry (GC-MS) analyses. Lipid peroxidation and cyclooxygenase enzyme inhibitory of these sterols were determined by using lipid peroxidation (LPO) and cyclooxygenase enzymes (COX-1 and −2) inhibitory assays according to our previous study.[16–21] Moreover, molecular docking analysis was performed to investigate the structure-activity relationships of these sterols.

## 2. Materials and methods

### 2.1 General experimental procedures

Waters 2010 HPLC (high pressure liquid chromatography) system (Waters Corp., Milford, MA, USA) equipped with XTerra Prep MS C-18 column (10 μm, 19 × 250 mm) (Waters Corporation, Milford, MA, USA) was used for the HPLC purification and all solvents used for HPLC purification were of ACS reagent grade (Sigma-Aldrich Chemical Company, St. Louis, MO, USA). The NMR (Nuclear magnetic resonance) spectra were obtained using Agilent DirectDrive2 500 MHz (Agilent Technologies, Palo Alto, CA, USA) in deuterated chloroform (CDCl_3_) purchased from Cambridge Isotope Laboratories, Inc. The ^1^H and ^13^C chemical shift values were expressed in ppm, based on the residual chemical shift values, for CDCl_3_ at *δ*H 7.24 and *δ*C 77.2 ppm, respectively. The COX-1 enzyme was prepared from ram seminal vesicles purchased from Oxford Biomedical Research, Inc. (Oxford, MI). The COX-2 enzyme was prepared from insect cells cloned with human PGHS-2 enzyme. Positive controls for the COX and LPO assays aspirin, naproxen, Celebrex^®^, ibuprofen, TBHQ (t-butylhydroquinone), BHA (butylated hydroxyanisole), BHT (butylated hydroxytoluene) and cholesterol were purchased from Sigma-Aldrich Chemical Company (St. Louis, MO, USA) and arachidonic acid was purchased from Oxford Biomedical Research, Inc (Oxford Biomedical Research, Inc., Oxford, MI, USA). The fluorescent probe, 3-[*p*-(6-phenyl)-1,3,5-hexatrienyl] phenylpropionic acid was purchased from Molecular Probes (Eugene, OR, USA) and 1-stearoyl-2-linoleoyl-sn-glycerol-3-phosphocholine (SLPC) was purchased from Avanti Polar Lipids (Alabaster, AL, USA).

### 2.2 Plant Materials

*L. japonica* (GS-02-004-2013) used in this study were cultivated at Yantai (Shandong province, China) and harvested in July 2018. Fresh raw *L. japonica* was dried immediately after harvest under sun light and then shipped to the lab. The raw *L. japonica* in the lab was first washed with running tap water and deionized water, and then dried in an oven at 50 °C. The dry sample was further pulverized with a blender to obtain the dry *L. japonica* power through a 50-mesh screen.

### 2.3 Extraction and isolation

The dry *L. japonica* power (5 kg) was extracted with Hexane (10 L, 12 h×3) and evaporated under vacuum to afford hexanes extract HE (22 g). An aliquot of HE (13 g) was pretreated with activated charcoal and then fractionated using silica gel VLC by eluting with hexanes-ethyl acetate (4:1, 2:1 and 1:1, v/v), followed by methanol (100%), to yield fractions A (855 mg), B (4 g), C (1.44 g), D (374 mg), E (205 mg), and F (3 g), respectively. An aliquot of fractions A (68 mg) was fractionated by Pre-TLC with hexanes-acetone (10: 1, *v/v*) to yield the main fraction A-1(31 mg). An aliquot of fractions B (76 mg) was fractionated by Pre-TLC with hexanes-acetone (8: 1, *v/v*) to yield the main fraction B-1(50 mg), which was characterized as fatty acid and hence were kept aside. Fraction D (374 mg) fractionated by MPLC with hexanes-acetone (2: 1, and 1: 1, *v/v*) and Pre-TLC with chloroform: methanol (40: 1, *v/v*) yielded three main fractions D-1(24 mg, 86B), D-2(2 mg, 91C) and D-3(20 mg, 72B), which were the same compounds that from fraction C. And fraction E and F were found to be salt and put away.

Fraction C (1.44 g) was further purified with silica gel MPLC by eluting with hexanes-acetone (4: 1, 2:1 and 1: 1, v/v), to yield fractions G (301 mg), H (599 mg), I (137 mg), J (160 mg), and K (166 mg), respectively. Fraction G had been put away for it was too complicated.

An aliquot of the main fraction **H** (215 mg) was fractionated by silica gel VLC and eluted under gradient conditions using chloroform: methanol (100:1 and 4.1 *v/v*) to yield fractions **LJ-1** (161 mg) and **LJ-2** (51 mg). An aliquot of fraction **LJ-1** (160 mg), purified by HPLC by eluting with acetonitrile: methanol 50:50 *v/v*, 4 mL/min under isocratic conditions yielded fraction **LJ-1(a)** (62 mg), compound **3**(39 mg, 42 min, 24-methylenecholesterol, **Figure S1 SI File**) and compound **4** (58 mg, 51 min, fucosterol, **Figure S1 SI File**).[22–24] Repeated HPLC purification of the fraction **LJ-1(a)** (40 mg) eluting with acetonitrile:methanol 95:5 *v/v*, 3 mL/min under isocratic conditions yielded compound **1** (12 mg, 46 min, 29-Hydroperoxy-stigmasta-5,24(28)-dien-3β-ol, **Figure S1 SI File**) and compound **2** (22 mg, 51 min, saringosterol **Figure S1 SI File**).[22]

Fraction I (137 mg) was purified by Prep-TLC on silica gel with chloroform-Methanol (40: 1) as the mobile phase, yielded fraction L (33 mg). An aliquot of fraction **L** (20 mg) was further separated by recrystallization from hexane: acetone to afford compound **5** (17 mg, 24-hydroperoxy-24-vinyl-cholesterol).[22]

Pre-TLC with chloroform: methanol (60: 1, *v/v*) of fraction J (51 mg) produced J-1 (15 mg, 87B) and J-2 (8 mg, 87D). Fraction K (75 mg) fractionated by Pre-TLC with hexanes-acetone (8: 1, *v/v*) afforded K-1 (43 mg, 69B). Fraction A-1, B-1, J-1, J-2, and K-1 mainly contained fatty acids and triglycerides based on preliminary analyses were kept aside due to prior published work. [25]

### 2.4 Cyclooxygenase enzymes (COX-1 and −2) inhibitory assays

The COX-1 and-2 enzyme inhibitory activity of pure isolates (compounds 1-5) from *L. japonica* and cholesterol were measured by monitoring the initial rate of O_2_ uptake using an Instech micro oxygen chamber and electrode (Instech Laboratories) attached to a YSI model 5300 biological oxygen monitor (Yellow Springs Instrument, Inc.) at 37 °C according to the published procedure.[25–28]

### 2.5 Lipid peroxidation inhibitory assay

The antioxidant activity of all isolates (compounds **1-5**) and cholesterol was determined by the LPO inhibitory assay using fluorescence spectroscopy on a Turner model 450 fluorometer (Barnstead/Thermolyne Corp.) according to the reported procedure.[17–20]

### 2.6 Molecular Docking

Docking was performed to investigate the molecular interactions between the pure compounds and cyclooxygenase using the AutoDock Vina open-source program (ver. 1.1.2) according to the reported procedure.[29, 30] The crystallographic structures of ovine COX-1 (PDB: 3N8Z) and human COX-2 (PDB: 5F1A) were retrieved from the Protein Data Bank in SDF format. The 3D conformations of compounds 1-5 were provided from PubChem (http://pubchem.ncbi.nlm.nih.gov/). Virtual screening of all compounds was performed into the active site of the COX-1 and COX-2 domain by using AutoDock Vina. The whole protein structures were targeted for compounds docking.[31] Molecular docking coordinates for COX-1 receptor were determined as center_x = −34.054, center_y =56.874, center_z = −11.092, dimensions parameters are size_x = 94, size_y = 96, size_z = 126, spacing parameter is 0.808 Å. Docking parameters used for COX-2 receptor were center_x = 34.099, center_y = 27.4, center_z = 217.829, size_x = 96, size_y = 96, size_z = 126, spacing parameter is 0.789 Å. Cluster analysis was performed on the docked results using an RMS tolerance of 2.0 Å. The docked formation with the lowest energy value was obtained, and enzyme-ligand interactions were visualized using open-source PyMOL created by Warren Lyford DeLano (https://github.com/schrodinger/pymol-open-source).

### 2.7 Statistical Analysis

The data was presented as mean ± SD (n = 3) and evaluated by one-way analysis of variance (ANOVA) followed by the Duncan’s test. All statistical analyses were carried out using R statistical package and SPSS for Windows, Version 17.0 (SPSS Inc., Chicago, IL, USA).

## 3. Results and discussion

### 3.1 Structure elucidation

Compound **1** was isolated as a white powder and exhibited a molecular ion peak at *m/z* 427.3563 [M-H_2_O+H]^+^ in its positive-ion HRESITOFMS consistent with the molecular formula C_29_H_48_O_3_, This confirmed six indices of hydrogen deficiency in compound **1** (**figure S2 SI File**). The ^1^H-NMR (500 MHz, CDCl_3_) spectrum exhibited two olefin methine proton signal at *δ*H 5.33 (1H, brd, J=4.9 Hz, H-6) and *δ*H 4.53 (1H, d, J=6.8 Hz, H-28), one oxygenated methine proton signal at *δ*H 3.49 (1H, m, H-3), and five methyl proton signals at *δ*H 0.99 (3H, s, H-19), 0.97 (3H, d, J=6.8 Hz, H-21), 0.95 (3H, d, J=6.8 Hz, H-27), 0.89 (3H, d, J=6.7 Hz, H-26), and 0.65 (3H, s, H-18) (**figure S3 SI File**). The ^13^C-NMR (125 MHz, CDCl_3_) spectrum exhibited 29 carbon signals including 2 olefin quaternary carbon signals at *δ*C 155.3 (C-24) and 140.7 (C-5), two olefin methine carbon signals at *δ*C121.7 (C-6) and *δ*C114.8 (C-28), one oxygenated methine carbon signal at *δ*C 71.8 (C-3), one oxygenated methylene carbon signal at *δ*C 73.5 (C-29) and 5 methyl carbon signals at *δ*C 21.9 (C-26), 21.8 (C-27), 19.4 (C-19), 18.7 (C-21), and 11.8 (C-18) (**figure S4 SI File**). According to the spectral data, compound **1** was identified as 29-Hydroperoxy-stigmasta-5,24(28)-dien-3β-ol (**Figure 1**).[22]

**Figure 1:**
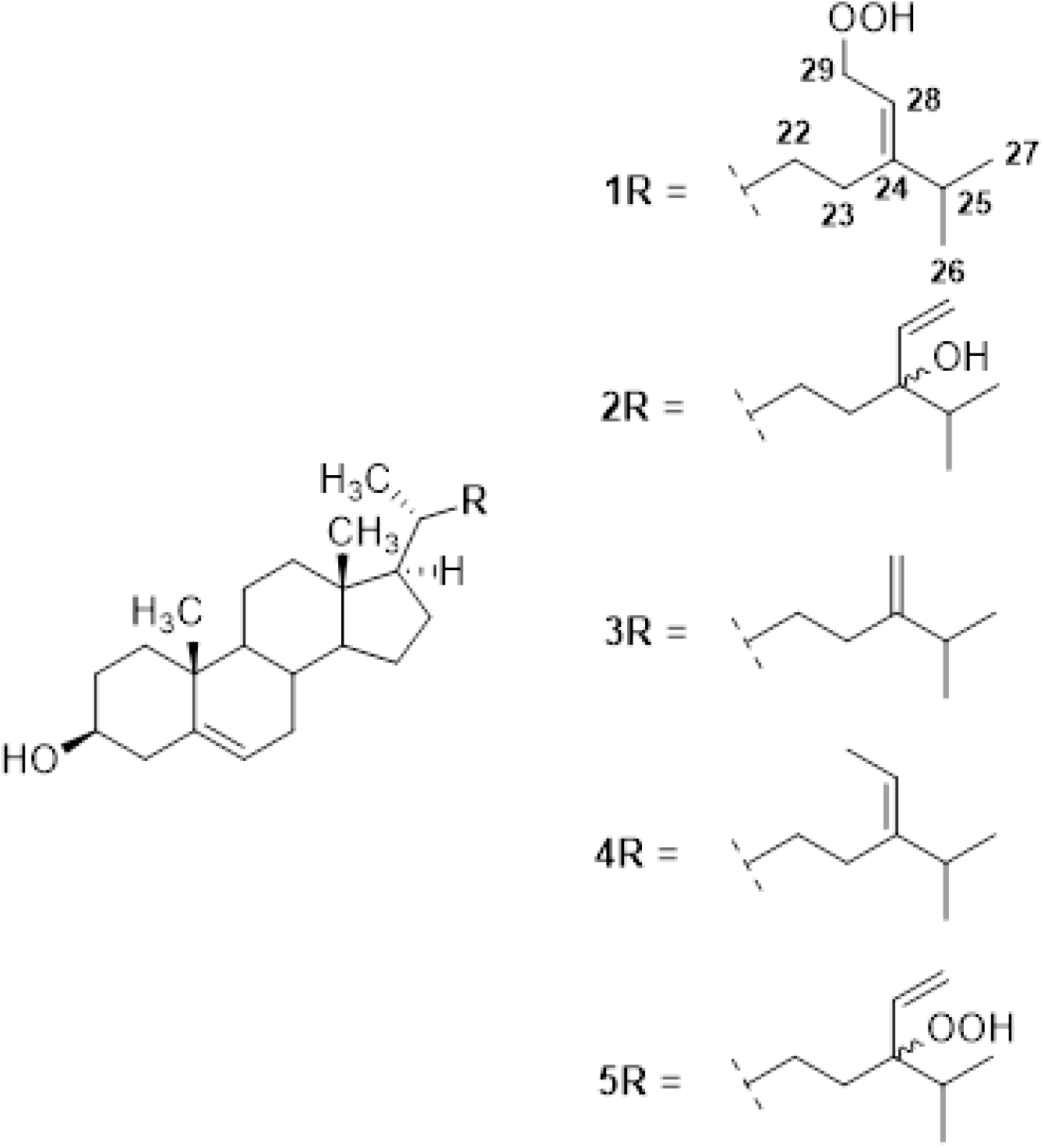
Chemical structures of the biologically active sterols (compounds **1-5**) isolated from the hexane extract of *L. japonica.*

Compound **2** was isolated as a white powder and exhibited a molecular ion peak at *m/z* 427.3562 [M+H-2H]^+^ in its positive-ion HRESITOFMS consistent with the molecular formula C_29_H_48_O_2_, This confirmed six indices of hydrogen deficiency in compound **2** (**figure S5 SI File**). The ^1^H-NMR (500 MHz, CDCl_3_) spectrum exhibited two olefin methine proton signals at δH 5.34 (1H, brd, J=4.9 Hz, H-6) and 5.75 (1H, m, H-28), one olefin methylene signal at δH 5.11 − 5.27 (2H, m, H-29a and H-29b), one oxygenated methine proton signal at δH 3.49 (1H, m, H-3), and five methyl proton signals at δH 0.98 (3H, s, H-19), 0.90 (3H, d, J=6.8 Hz, H-21), 0.89 (3H, d, J=6.8 Hz, H-27), 0.85 (3H, d, J=6.7 Hz, H-26), and 0.66 (3H, s, H-18) (**figure S6 SI File**). The ^13^C-NMR (125 MHz, CDCl_3_) spectrum exhibited 29 carbon signals including one olefin quaternary carbon signal at δC 140.7 (C-5), one olefin methylene carbon signals at *δ*C121.7 (C-29), two olefinic methines carbon signals at *δ*C137.1 (C-5) and *δ*C116.3 (C-28), two oxygenated methine carbon signal at δC 71.7 (C-3) and δC 89.1 (C-24), and five methyl carbon signals at δC 21.0 (C-26), 19.3 (C-27), 18.7 (C-19), 17.5 (C-21), and 11.8 (C-18) (**figure S7 SI File**). Therefore, according to the spectral data, compound **2** was identified as saringosterol (24-vinyl-cholest-5-ene-3β,24-diol) (**Figure 1**). [22]

Compound **3** was isolated as a white powder and exhibited a molecular ion peak at *m/z* 381.3507 [M-H_2_O+H]^+^ in its positive-ion HRESITOFMS consistent with the molecular formula C_28_H_46_O, This confirmed six indices of hydrogen deficiency in compound **3** (**figure S8 SI File**). The ^1^H-NMR (500 MHz, CDCl_3_) spectrum exhibited one olefin methine proton signal at *δ*H 5.33 (1H, brd, J=4.9 Hz, H-6), one olefinic methylene proton signals at *δ*H 4.69 (1H, d, J=1.0, H-28a) and 4.64 (1H, d, J=1.0, H-28b), one oxygenated methine proton signal at *δ*H 3.49 (1H, m, H-3), and five methyl proton signals at *δ*H 0.98 (3H, s, H-19), 0.97 (3H, d, J=6.8 Hz, H-21), 0.96 (3H, d, J=6.8 Hz, H-27), 0.94 (3H, d, J=6.7 Hz, H-26), and 0.66 (3H, s, H-18) (**figure S9 SI File**). The ^13^C-NMR (125 MHz, CDCl_3_) spectrum exhibited 28 carbon signals including 2 olefin quaternary carbon signals at *δ*C 156.9 (C-24) and 140.7 (C-5), one olefin methine carbon signals at *δ*C121.7 (C-6), one olefin methylene carbon signals at *δ*C105.9 (C-28), one oxygenated methine carbon signal at *δ*C 71.8 (C-3), and 5 methyl carbon signals at *δ*C 21.9 (C-26), 21.8 (C-27), 19.4 (C-19), 18.7 (C-21), and 11.8 (C-18) (**figure S10 SI File**). According to the spectral data, compound **3** was identified as 24-methylenecholesterol (**Figure 1**).[22, 23]

Compound **4** was also isolated as a white powder and exhibited a molecular ion peak at *m/z* 395.3669 [M-H_2_O+H]^+^ in its positive-ion HRESITOFMS consistent with the molecular formula C_29_H_48_O, This confirmed six indices of hydrogen deficiency in compound **4** (**figure S11 SI File**). The ^1^H-NMR (500 MHz, CDCl_3_) spectrum exhibited two olefin methine proton signals at *δ*H 5.33 (1H, brd, J=4.9 Hz, H-6) and 5.16 (1H, dd, J=6.8 and 6.8 Hz, H-28), one oxygenated methine proton signal at *δ*H 3.49 (1H, m, H-3), one allyl methyl proton signal at *δ*H 1.55 (3H, d, J=6.8 Hz, H-29), and five methyl proton signals at *δ*H 0.98 (3H, s, H-19), 0.97 (3H, d, J=6.8 Hz, H-21), 0.96 (3H, d, J=6.8 Hz, H-27), 0.94 (3H, d, J=6.7 Hz, H-26), and 0.66 (3H, s, H-18) (**figure S12 SI File**). The ^13^C-NMR (125 MHz, CDCl_3_) spectrum exhibited 29 carbon signals including 2 olefin quaternary carbon signals at *δ*C 146.9 (C-24) and 140.7 (C-5), two olefin methine carbon signals at *δ*C121.7 (C-6) and 115.5 (C-28), one oxygenated methine carbon signal at *δ*C 71.7 (C-3), and six methyl carbon signals at *δ*C 13.2 (C-29), 22.2 (C-26), 22.1 (C-27), 19.4 (C-19), 18.7 (C-21), and 11.8 (C-18) (**figure S13 SI File**). Therefore, according to the spectral data, compound **4** was identified as fucosterol (stigmasta-5,24-diene-3β-ol) (**Figure 1**).[22, 26]

Compound **5** was also isolated as a white powder and exhibited a molecular ion peak at *m/z* 427.3562 [M-H_2_O+H]^+^ in its positive-ion HRESITOFMS consistent with the molecular formula C_29_H_48_O_3_, This confirmed six indices of hydrogen deficiency in compound **5** (**figure S14 SI File**). The ^1^H-NMR (500 MHz, CDCl_3_) spectrum exhibited two olefin methine proton signals at δH 5.34 (1H, brd, J=4.9 Hz, H-6) and 5.75 (1H, m, H-28), one olefin methylene signal at δH 5.09 - 5.19 (2H, m, H-29a and H-29b), one oxygenated methine proton signal at δH 3.49 (1H, m, H-3), and five methyl proton signals at δH 0.99 (3H, s, H-19), 0.91 (3H, d, J=6.8 Hz, H-21), 0.89 (3H, d, J=6.8 Hz, H-27), 0.86 (3H, d, J=6.7 Hz, H-26), and 0.64 (3H, s, H-18) (**figure S15 SI File**). The ^13^C-NMR (125 MHz, CDCl_3_) spectrum exhibited 29 carbon signals including one olefin quaternary carbon signal at δC 140.7 (C-5), one olefin methylene carbon signals at *δ*C121.6 (C-29), two olefinic methines carbon signals at *δ*C 112.8 (C-5) and *δ*C 142.5 (C-28), two oxygenated methine carbon signal at δC 71.7 (C-3) and δC 77.7 (C-24), and five methyl carbon signals at δC 21.0 (C-26), 19.3 (C-27), 18.7 (C-19), 17.5 (C-21), and 11.8 (C-18) (**figure S16 SI File**). Therefore, according to the spectral data, compound **5** was identified as 24-Hydroperoxy-24-vinyl-cholesterol (**Figure 1**).[22]

### 3.2 Cyclooxygenase enzymes (COX-1 and −2) inhibitory assay

Prostaglandins, the inflammation-causing hormones, were converted from arachidonic acid by the catalysis of COX enzymes. Therefore, inhibition of COX enzymes could prevent the production of prostaglandins and reduce inflammation.[26, 32] The anti-inflammatory activity of the pure isolates from *L. japonica* was revealed by their COX-1 and −2 enzyme inhibitions. In this study, compounds (**1-5**) at 25 μg/mL inhibited COX-1 enzyme by 32, 19, 50, 59, and 26% and COX-2 enzyme by 23, 14, 33, 47, and 9%, respectively (**Figure 2**). Commercial NSAIDs aspirin, Celebrex^®^, naproxen and ibuprofen were used as positive control at 108, 1, 12, and 15 μg/mL, respectively (**Figure 2**). Apparently, the COX enzyme inhibitory activity of 24-methylenecholesterol (compound 3) and fucosterol (compound 4) was comparable to the activity of the over-the-counter nonsteroidal anti-inflammatory drugs (NSAIDs) aspirin and ibuprofen. Compound 3 and 4 inhibited the COX-1 enzyme at a higher rate than the COX-2 enzyme, which is similar to aspirin and ibuprofen.

**Figure 2:**
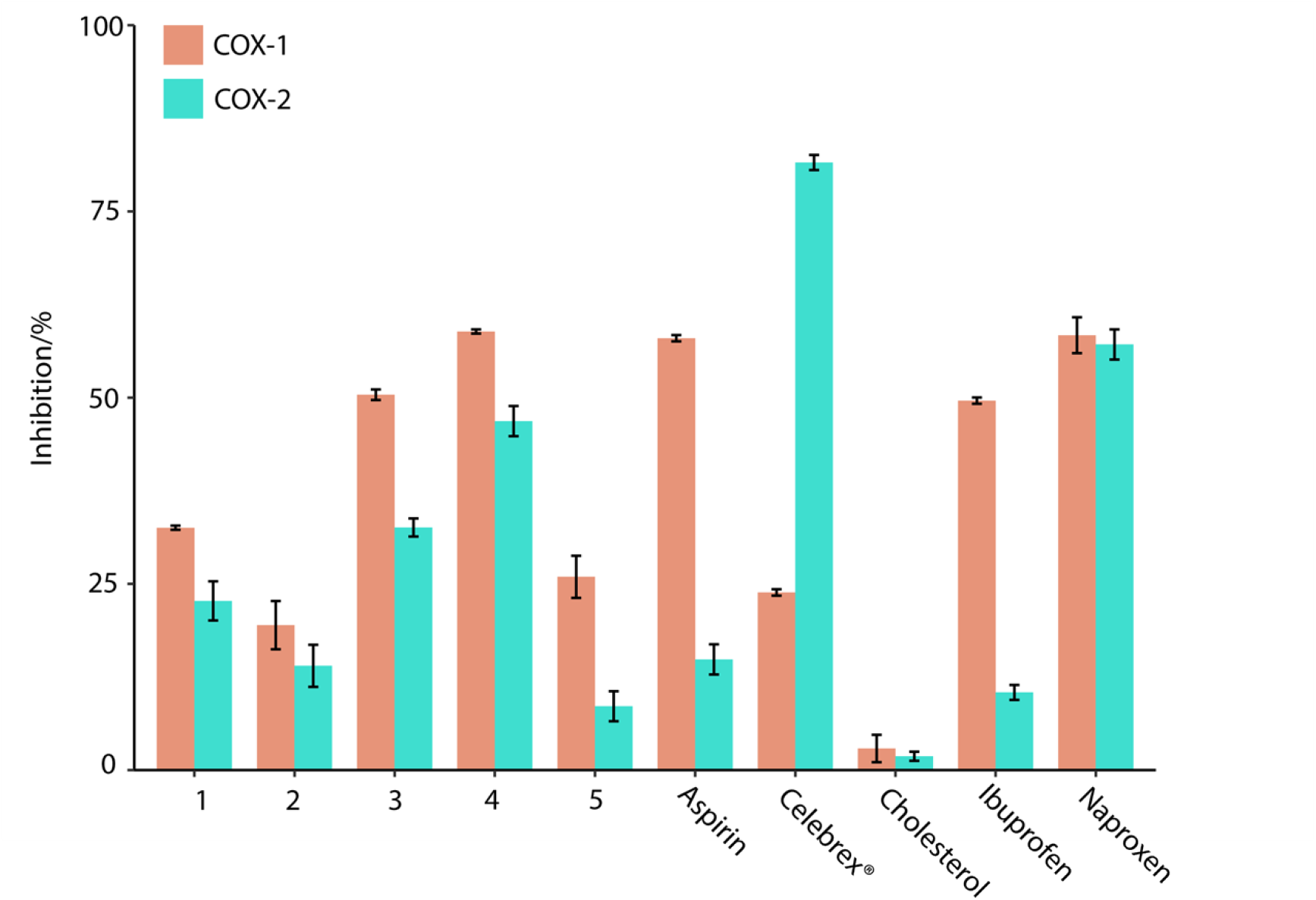
Cyclooxygenase enzymes (COX-1 and −2) enzyme inhibitory activities sterols (**1-5**) from the hexane extract of *L. japonica* and cholesterol were tested at 25 μg/mL concentration. Commercial NSAIDs aspirin, ibuprofen, Celebrex^®^ and naproxen used as positive control and tested at 108, 12, 1 and 12 μg/mL, respectively. The varying concentrations of positive controls used were to yield a comparable activity profile between 50-100% by test compounds and positive controls alike. Vertical bars represent the standard deviation of each data point (n=2).

### 3.3 Lipid peroxidation inhibitory assay

Potential antioxidant activity of the pure compounds 1-5 was determined by the LPO assays using the large unilamellar vesicles (LUVs) model system, the results of which are shown in Fig. 3. The LPO assay detects compounds that can scavenge free radicals. At 25 μg/mL, the LPO inhibitory activity of sterols **1-5** and cholesterol was 21, 51, 58, 56, 25 and 20%, respectively, as compared to the commercial antioxidants BHA, BHT and TBHQ at 85, 85 and 82%, respectively. The concentrations used to test LPO inhibitory activity for compounds were at 1.8, 2.2 and 1.6 μg/mL, respectively (**Figure 3**). Compared to compound 1 and 5, saringosterol (compound 2), 24-methylenecholesterol (compound 3) and fucosterol (compound 4) showed the higher LPO inhibitory activity at >50%, at 25 μg/mL concentration.

**Figure 3:**
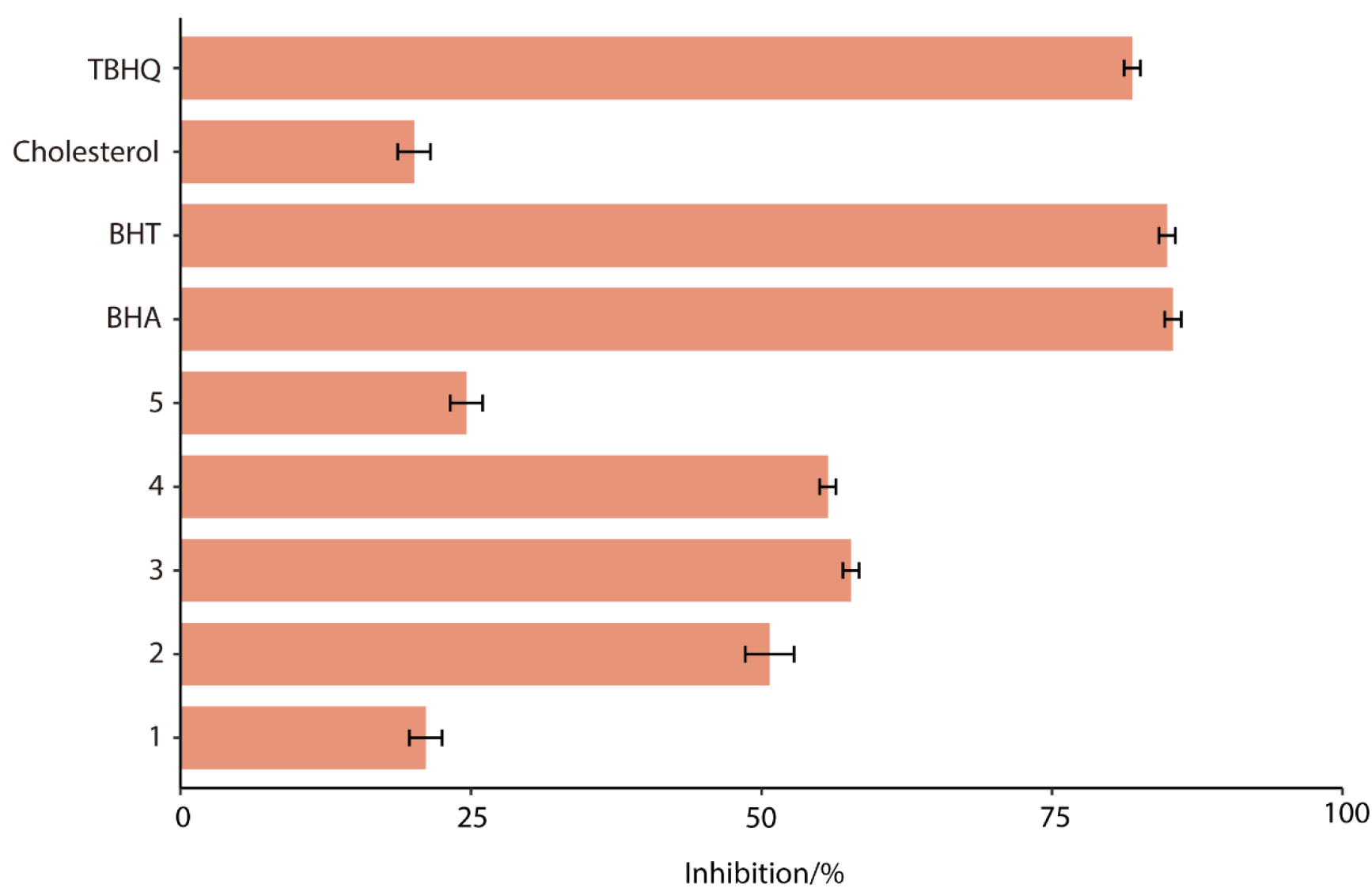
Lipid peroxidation (LPO) inhibitory activities of sterols (**1-5**) isolated from the hexane extract of *L. japonica* and commercial cholesterol tested at 25 μg/mL. Commercial antioxidants BHA, BHT and TBHQ used as positive controls at 1.8, 2.2 and 1.6 μg/mL. The oxidation of lipid was initiated by the addition of Fe^2+^ ions. The varying concentrations of positive controls used were to yield a comparable activity profiles between 50-100% by test compounds and positive controls alike. Vertical bars represent the standard deviation of each data point (n=2).

### 3.4 Molecular docking

Molecular docking was used to predict the interactions between the 5 sterols and the COX-1 and −2 enzyme at the molecular level. Root-Mean-Squared-Deviation (RMSD) is a similarity metric between molecular conformations, the lower the RMSD value is, the more similar the molecular conformations of ligands (before and after the docking) is. As shown in Table 1, the RMSD values of all the detected ligands were below the maximum allowed value of 2 Å, indicating ligands underwent minimal deformation during docking and the docking results were credible.[33] Docking results revealed that the binding energy of compounds 1-5 were −7.64, −6.36, −7.79, −7.85 and −6.44 kcal/mol for COX-1, and −8.49, −8.33, −8.72, −9.02 and −6.97 kcal/mol for COX-2, which have the similar trends with their COX-1 and −2 enzyme inhibitory activities (Table 1).

**Table 1.** Docking results of all the 5 sterols against COX-1 and −2 receptors.

| Receptor<br>with PDB<br>IDs | Ligand | Binding<br>affinity<br>(kcal/mol) | RMSD(Å) | No. of H-bonds | Amino acid<br>residues forming<br>H-bonds |
| --- | --- | --- | --- | --- | --- |
| COX-1<br>(3N8Z) | Comp. 1 | -7.64 | 1.28 | 3 | ASP-135,<br>ASP-158 |
|  | Comp. 2 | -6.36 | 1.43 | 1 | GLU-326 |
|  | Comp. 3 | -7.79 | 1.09 | 1 | ASP-158 |
|  | Comp. 4 | -7.85 | 1.08 | 0 | None |
|  | Comp. 5 | -6.44 | 1.69 | 1 | LEU-115 |
| COX-2<br>(5F1A) | Comp. 1 | -8.49 | 0.88 | 2 | GLY-225,<br>TYR-373 |
|  | Comp. 2 | -8.33 | 1.82 | 2 | ASN-375 |
|  | Comp. 3 | -8.72 | 1.06 | 1 | GLY-225 |
|  | Comp. 4 | -9.02 | 1.43 | 1 | GLY-225 |
|  | Comp. 5 | -6.97 | 1.83 | 2 | ARG-120,<br>GLU-524 |

The possible mechanisms beneath the inhibition of compounds 1-5 against COX-1 and −2 was presented through molecular docking (Fig. 4–6, Table 1). As shown in Fig. 4, compounds 1-5 were predicted to be located in different pockets of COX-1 (pocket A, B and C), as reported in previous studies. [34] The hydrogen bonds and interaction residues were displayed in 3D docked poses of compounds 1-5 binding to COX-1 and COX-2 (Fig. 6). Compounds 1 and 4 located in the same active pocket of COX-1, which was surrounded by active site at ILE-46, CYS-47, CYS-36, ASN-34, ARG-49, THR-322, GLY-324, GLU-326, GLN-327, TYR-136, ASP-135 and PRO-153. The docking results also revealed that compound 1 typically bind within the COX-1 channel via H-bonding interactions with the side chain of ASP-135 and ASP-158, while compound 4 was embedded in the active pocket of COX-1 primarily through hydrophobic interactions (Fig.6, Table 1).

**Figure 4:**
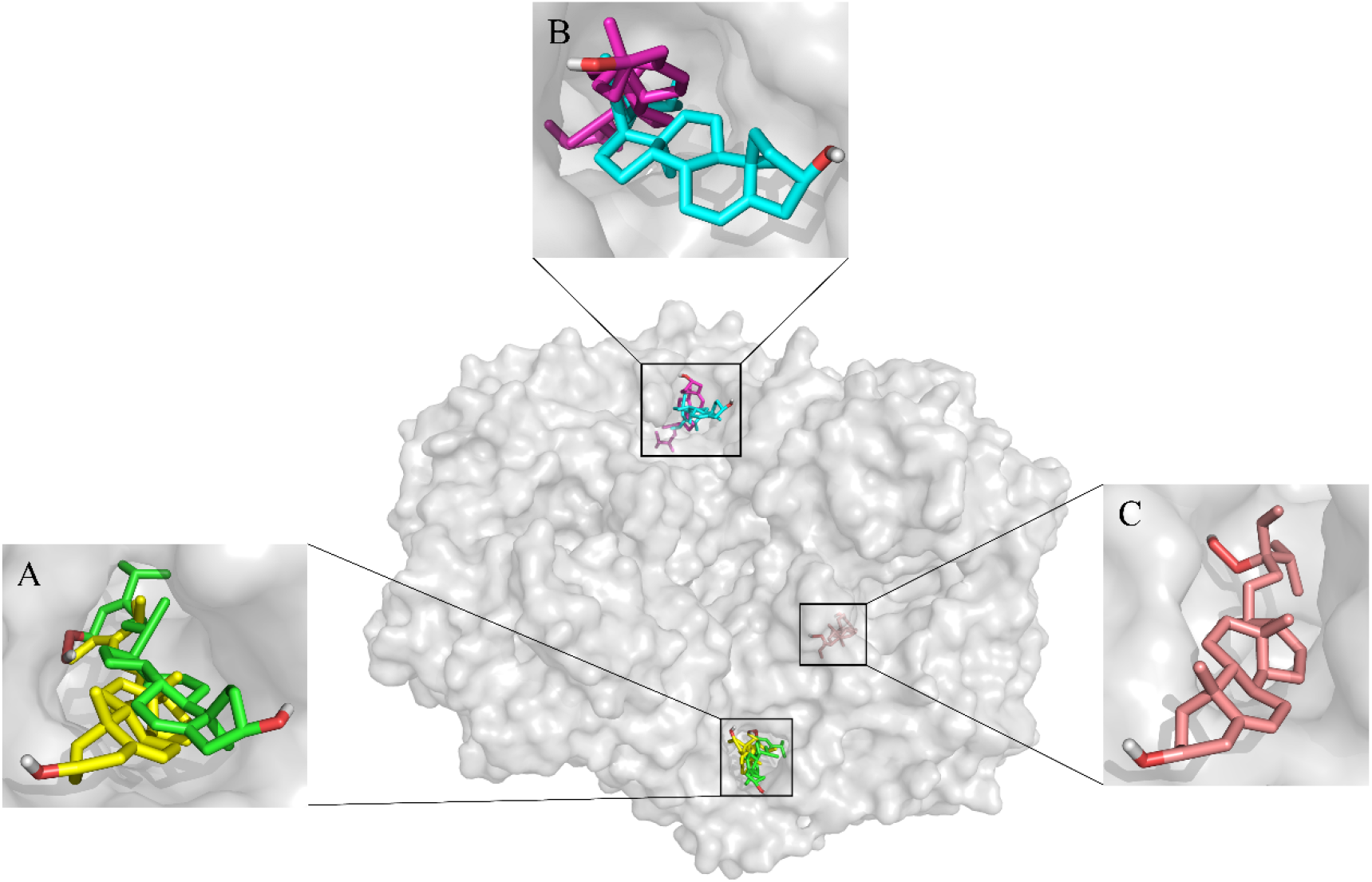
Molecular docking of COX-1 with compounds 1-5. Different compounds were located in different active pockets. Compounds 1 and 4 were located in the hydrophobic pocket A of COX-1 (Comp. 1 shown in green, Comp.4 shown in yellow), compounds 2 and 3 were located in the pocket B of COX-1 (Comp.2 shown in cyan, Comp.3 shown in magenta), while compound 5 was located in the pocket C (Comp.5 shown in salmon).

**Figure 5:**
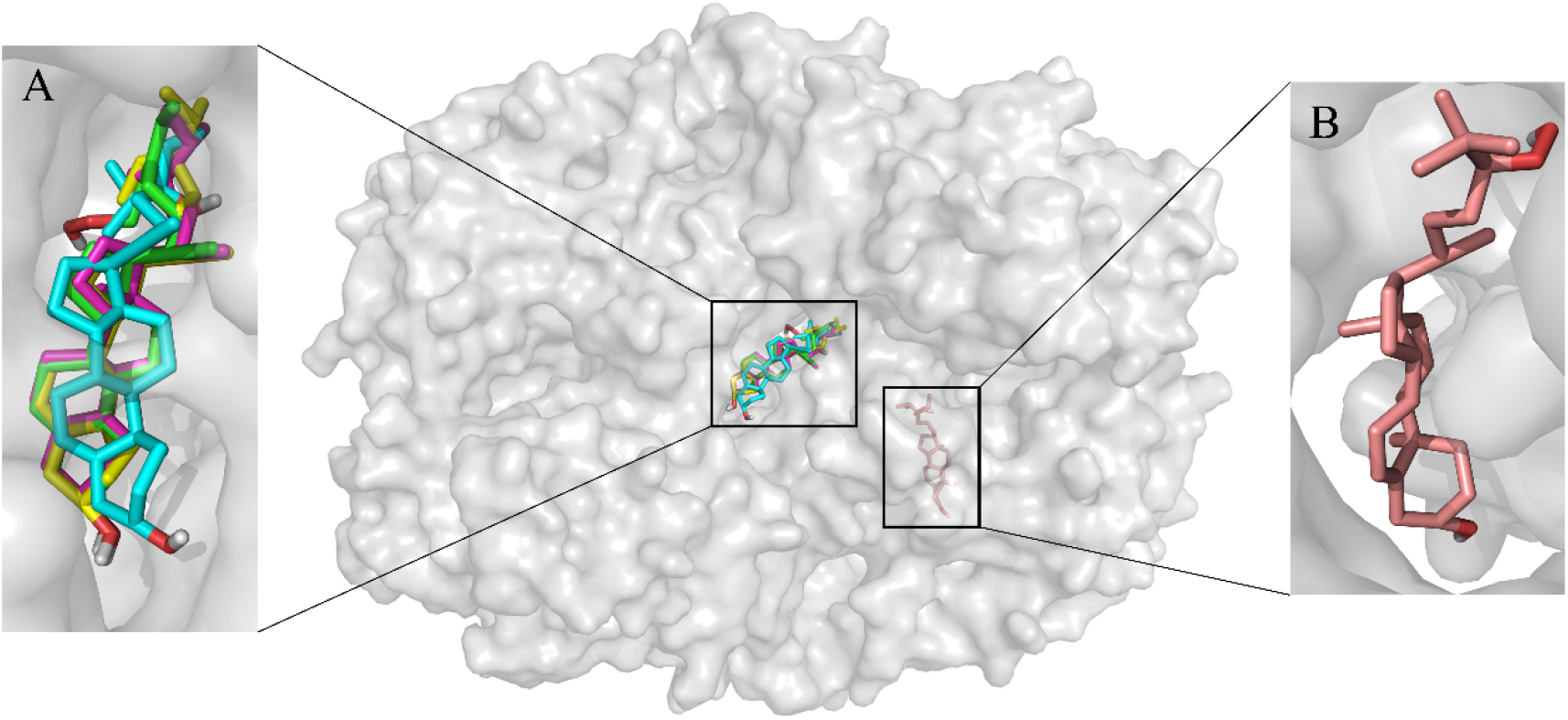
Molecular docking of COX-2 with compounds 1-5. Different compounds were located in different active pockets. Compounds 1, 2, 3 and 4 were located in the hydrophobic pocket A of COX-2 (Comp. 1 shown in green, Comp.2 shown in cyan, Comp.3 shown in magenta, Comp.4 shown in yellow), while compound 5 was located in the pocket B (Comp.5 shown in salmon).

**Figure 6:**
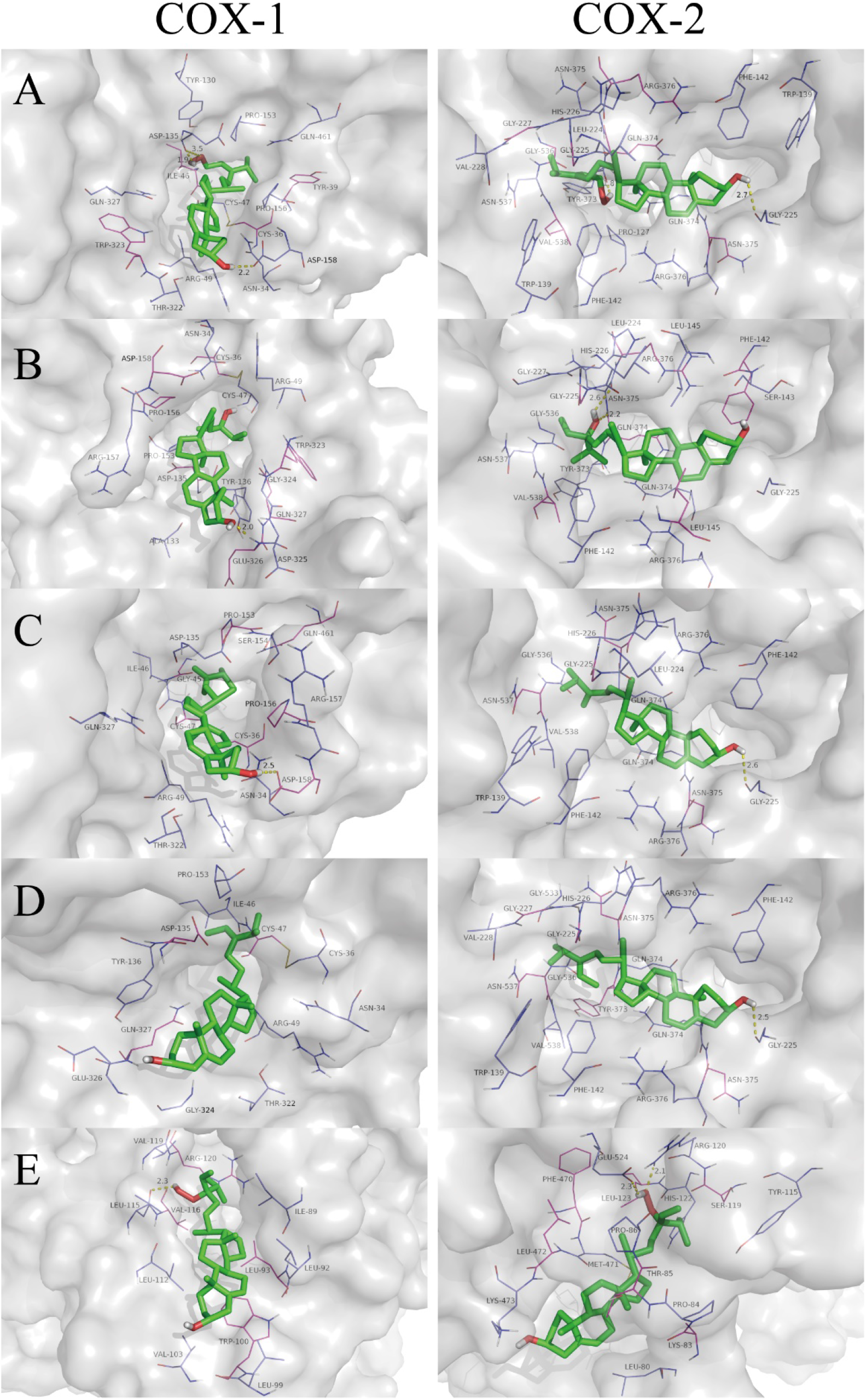
3-D docked poses of COX-1 and −2 with five sterols (compounds 1-5). The first column represents COX-1 and the second column represents COX-2. The rows present: A) compound 1, B) compound 2, C) compound 3, D) compound 4, E) compound 5.

Furthermore, the binding sites and interactions between compounds 1-5 and COX-2 are represented in Fig 5 and 6. Compounds 1-4 were located in the hydrophobic pocket A of COX-2 (surrounded by active site at GLY-225, ASN-375, ARG-376, PHE-142, VAL-538, ASN-537, GLY-227, GLY-536, LEU-224, HIS-226, GLN-374, TYR-373), while compound 5 was located in the pocket B (surrounded by active site at PRO-84, LYS-83, LEU-80, LYS-473, LEU-472, PHE-470, GLU-524, ARG-120, TYR-115, THR-85, MET-471, PRO-86, LEU-123, HIS-122, SER-119). All of the five compounds had interactions with surrounding residues in the active site of COX-2. Compounds 1, 3 and 4 showed possible hydrogen bond with GLY-225, predicting which might be favorable binding site of COX-2 for sterols (Fig.6). Differ from compounds 1-4, compound 5 bound with COX-2 at another active pocket and showed possible hydrogen bonds with ARG-120 and GLU-524. Moreover, there was a high binding energy between compound 4 and COX-2 (−9.02 kcal/mol), which was comparable to that of the compound 5 (−6.97 kcal/mol).

COX-1 and −2 are bifunctional enzymes that convert arachidonic acid (AA) to prostaglandin G_2_ (PGG_2_) in their cyclooxygenase active site.[35] Our results suggested that fucosterol (compound 4) was located in the hydrophobic pocket of COX-1 and −2 enzyme with the highest binding energy (−7.85 and −9.02 kcal/mol) in molecular docking, compared to the other sterols. The possible mechanism underlying the highest COX-1 and −2 enzyme inhibitory activities of fucosterol might be attributed to the distinct olefin methine presented in its molecular structure, which could occupy the active site of the enzyme and prevent the enzyme from contacting with the substrate. This is in agreement with previous molecular docking results that tetracyclic skeletons and incorporation of an aliphatic chain in sterols were the key structural requirements theoretically for their good COX-1 and COX-2 binding activity.[36] Thus, these findings confirmed that fucosterol with tetracyclic skeletons and olefin methine achieved the good COX-1 and −2 enzyme inhibitory activities through hydrophobic interactions and hydrogen bond.

## 4. Conclusions

Purification of hexane extracts from *L. japonica* afforded 5 sterols, and their structures were identified using spectroscopic and chemical evidence. Spectroscopic methods characterized these compounds as 29-Hydroperoxy-stigmasta-5,24(28)-dien-3β-ol (compound 1), saringosterol (24-vinyl-cholest-5-ene-3β,24-diol) (compound 2), 24-methylenecholesterol (compound 3), fucosterol (stigmasta-5,24-diene-3β-ol) (compound 4), and 24-Hydroperoxy-24-vinyl-cholesterol (compound 5). These pure isolates were tested for antioxidant and anti-inflammatory activities using LPO and COX-1 and −2 enzyme inhibitory assays. Compared to the other compounds, fucosterol (compound 4) exhibited the highest COX-1 and −2 enzyme inhibitory activities at 59 and 47%, respectively. The COX enzyme inhibitory activity of 24-methylenecholesterol (compound 3) and fucosterol (compound 4) was comparable to the activity of the NSAIDs aspirin and ibuprofen. For the LPO assays, saringosterol (compound 2), 24-methylenecholesterol (compound 3) and fucosterol (compound 4) showed higher LPO inhibitory activity at >50% than the other compounds. In addition, the results of molecular docking predicted that 5 sterols were located in different pocket of COX-1 and −2 and confirmed that fucosterol with tetracyclic skeletons and olefin methine achieved the best COX-1 and −2 enzyme inhibitory activities through hydrophobic interactions and hydrogen bond. Our results confirm the presence of 5 sterols in *L. japonica* and its significant anti-inflammatory and antioxidant activity. Therefore, it appears that *L. japonica* have the potential to be developed as a dietary supplement.

## Abbreviations

HRESITOFMS: high-resolution electrospray ionization mass spectrometry
NMR: nuclear magnetic resonance
LPO: lipid peroxidation
COX: cyclooxygenase
MPLC: medium-pressure liquid chromatography
VLC: vacuum liquid chromatography
Prep-TLC: preparative thin-layer chromatography
HPLC: high pressure liquid chromatography
TBHQ: t-butylhydroquinone
BHA: butylated hydroxyanisole
BHT: butylated hydroxytoluene
NSAIDs: nonsteroidal anti-inflammatory drugs
SLPC: 1-stearoyl-2-linoleoyl-sn-glycerol-3-phosphocholine
LUVs: large unilamellar vesicles
AA: arachidonic acid

## Acknowledgement

Jie Gao gratefully acknowledge the financial support from China Scholarship Council (CSC) and National Natural Science Foundation of China (Project No. 32001696). This research is also a contribution from Michigan State University AgBioResearch (MICL01680). The nuclear magnetic resonance data were obtained on instrumentation that was purchased, in part, with the funds from NIH grant no. 1-S10-RR04750, NSF grant no. CHE-88-00770, and NSF grant no. CHE-92-13241. HRMS data were obtained at the Michigan State University Mass Spectrometry Facility, which is supported, in part, by a grant (DRR-00480) from the Biotechnology Research Technology Program, National Centre for Research Resources, National Institutes of Health.

## Author Contributions

Muraleedharan G. Nair and Mouming Zhao conceived the idea. Xingyu Lu and Jie Gao conducted the experiment and write manuscript. Amila Dissanayake analyzed the data. Chuqiao Xiao conducted the molecular docking. All authors contributed to the interpretation and discussion of the results.

## References

1. Zha X-Q, Xiao J-J, Zhang H-N, Wang J-H, Pan L-H, Yang X-F, et al. Polysaccharides in Laminaria japonica (LP): Extraction, physicochemical properties and their hypolipidemic activities in diet-induced mouse model of atherosclerosis. Food Chemistry. 2012;134(1):244–52.

2. Ren R, Azuma Y, Ojima T, Hashimoto T, Mizuno M, Nishitani Y, et al. Modulation of platelet aggregation-related eicosanoid production by dietary F-fucoidan from brown alga Laminaria japonica in human subjects. British journal of nutrition. 2013;110(5):880–90.

3. Gao J, Lin L, Chen Z, Cai Y, Xiao C, Zhou F, et al. In Vitro Digestion and Fermentation of Three Polysaccharide Fractions from Laminaria japonica and Their Impact on Lipid Metabolism-Associated Human Gut Microbiota. Journal of Agricultural and Food Chemistry. 2019;67(26):7496–505. doi: 10.1021/acs.jafc.9b00970.

4. Zhao D, Xu J, Xu X. Bioactivity of fucoidan extracted from Laminaria japonica using a novel procedure with high yield. Food Chemistry. 2018;245:911–8. doi: https://doi.org/10.1016/j.foodchem.2017.11.083.

5. Han D, Zhu T, Row KH. Ultrasonic extraction of phenolic compounds from Laminaria japonica Aresch using ionic liquid as extraction solvent. Bulletin of the Korean Chemical Society. 2011;32(7):2213.

6. Liu C-L, Chiu Y-T, Hu M-L. Fucoxanthin enhances HO-1 and NQO1 expression in murine hepatic BNL CL. 2 cells through activation of the Nrf2/ARE system partially by its pro-oxidant activity. Journal of agricultural and food chemistry. 2011;59(20):11344–51.

7. Liu Y, Liu M, Zhang X, Chen Q, Chen H, Sun L, et al. Protective effect of fucoxanthin isolated from Laminaria japonica against visible light-induced retinal damage both in vitro and in vivo. Journal of agricultural and food chemistry. 2016;64(2):416–24.

8. Heo S-J, Yoon W-J, Kim K-N, Ahn G-N, Kang S-M, Kang D-H, et al. Evaluation of anti-inflammatory effect of fucoxanthin isolated from brown algae in lipopolysaccharide-stimulated RAW 264.7 macrophages. Food and Chemical Toxicology. 2010;48(8-9):2045–51.

9. Woo MN, Jeon SM, Shin YC, Lee MK, Kang MA, Choi MS. Anti‐obese property of fucoxanthin is partly mediated by altering lipid‐regulating enzymes and uncoupling proteins of visceral adipose tissue in mice. Molecular Nutrition & Food Research. 2009;53(12):1603–11.

10. Liu M, Zhang W, Qiu L, Lin X. Synthesis of butyl-isobutyl-phthalate and its interaction with α-glucosidase in vitro. Journal of Biochemistry. 2011;149(1):27–33.

11. Bu T, Liu M, Zheng L, Guo Y, Lin X. α‐glucosidase inhibition and the in vivo hypoglycemic effect of butyl ‐ isobutyl ‐ phthalate derived from the Laminaria japonica rhizoid. Phytotherapy Research. 2010;24(11):1588–91.

12. Wang C, Yang Y, Mei Z, Yang X. Cytotoxic compounds from Laminaria japonica. Chemistry of Natural Compounds. 2013;49(4):699–701.

13. Nishizawa M, Takahashi N, Shimozawa K, Aoyama T, Jinbow K, Noguchi Y, et al. Cytotoxic constituents in the holdfast of cultivated Laminaria japonica. Fisheries science. 2003;69(3):639–43.

14. Hussain T, Tan B, Yin Y, Blachier F, Tossou MC, Rahu N. Oxidative stress and inflammation: what polyphenols can do for us? Oxidative medicine and cellular longevity. 2016;2016.

15. Lee J-Y, Lee M-S, Choi H-J, Choi J-W, Shin T, Woo H-C, et al. Hexane fraction from Laminaria japonica exerts anti-inflammatory effects on lipopolysaccharide-stimulated RAW 264.7 macrophages via inhibiting NF-kappaB pathway. European journal of nutrition. 2013;52(1):409–21.

16. Liu Y, Singh D, Nair MG. Pods of Khejri (Prosopis cineraria) consumed as a vegetable showed functional food properties. Journal of Functional Foods. 2012;4(1):116–21.

17. Arora A, Nair MG, Strasburg GM. Antioxidant activities of isoflavones and their biological metabolites in a liposomal system. Archives of biochemistry and biophysics. 1998;356(2):133–41.

18. Dissanayake AA, Zhang C-R, Mills GL, Nair MG. Cultivated maitake mushroom demonstrated functional food quality as determined by in vitro bioassays. Journal of Functional Foods. 2018;44:79–85.

19. Dissanayake AA, Ameen BA, Nair MG. Lipid peroxidation and cyclooxygenase enzyme inhibitory compounds from Prangos haussknechtii. Journal of natural products. 2017;80(9):2472–7.

20. Zhang C-R, Dissanayake AA, Nair MG. Functional food property of honey locust (Gleditsia triacanthos) flowers. Journal of Functional Foods. 2015;18:266–74.

21. Jayaprakasam B, Alexander-Lindo RL, DeWitt DL, Nair MG. Terpenoids from Stinking toe (Hymneae courbaril) fruits with cyclooxygenase and lipid peroxidation inhibitory activities. Food Chemistry. 2007;105(2):485–90. doi: https://doi.org/10.1016/j.foodchem.2007.04.004.

22. Chen Z, Liu J, Fu Z, Ye C, Zhang R, Song Y, et al. 24 (S)-Saringosterol from edible marine seaweed Sargassum fusiforme is a novel selective LXRβ agonist. Journal of agricultural and food chemistry. 2014;62(26):6130–7.

23. Huh G-W, Lee D-Y, In S-J, Lee D-G, Park SY, Yi T-H, et al. Fucosterols from Hizikia fusiformis and their proliferation activities on osteosarcoma-derived cell MG63. Journal of the Korean Society for Applied Biological Chemistry. 2012;55(4):551–5.

24. Hwang SH, Jang JM, Lim SS. Isolation of fucosterol from Pelvetia siliquosa by high-speed countercurrent chromatography. Fisheries and aquatic sciences. 2012;15(3):191–5.

25. Liu Y, Kakani R, Nair MG. Compounds in functional food fenugreek spice exhibit anti-inflammatory and antioxidant activities. Food Chemistry. 2012;131(4):1187–92. doi: https://doi.org/10.1016/j.foodchem.2011.09.102.

26. Kelm M, Nair M, Strasburg G, DeWitt D. Antioxidant and cyclooxygenase inhibitory phenolic compounds from Ocimum sanctum Linn. Phytomedicine. 2000;7(1):7–13.

27. Aoyagi M, Imai S, Kamoi T. Novel bisthiolane polysulfides from lachrymatory factor synthase-suppressed onion and their in vitro cyclooxygenase-1 inhibitory activity. Food Chemistry. 2021;344:128636. doi: https://doi.org/10.1016/j.foodchem.2020.128636.

28. Grancieri M, Martino HSD, Gonzalez de Mejia E. Digested total protein and protein fractions from chia seed (Salvia hispanica L.) had high scavenging capacity and inhibited 5-LOX, COX-1-2, and iNOS enzymes. Food Chemistry. 2019;289:204–14. doi: https://doi.org/10.1016/j.foodchem.2019.03.036.

29. Trott O, Olson AJ. AutoDock Vina: improving the speed and accuracy of docking with a new scoring function, efficient optimization, and multithreading. Journal of computational chemistry. 2010;31(2):455–61.

30. Huang R, Zhang Y, Shen S, Zhi Z, Cheng H, Chen S, et al. Antioxidant and pancreatic lipase inhibitory effects of flavonoids from different citrus peel extracts: An in vitro study. Food Chemistry. 2020;326:126785. doi: https://doi.org/10.1016/j.foodchem.2020.126785.

31. Vidal PJ, Lopez-Nicolas JM, Gandia-Herrero F, Garcia-Carmona F. Inactivation of lipoxygenase and cyclooxygenase by natural betalains and semi-synthetic analogues. Food Chem. 2014;154:246–54. Epub 2014/02/13. doi: 10.1016/j.foodchem.2014.01.014. PubMed PMID: 24518339.

32. Zhang C-R, Aldosari SA, Vidyasagar PS, Nair KM, Nair MG. Antioxidant and anti-inflammatory assays confirm bioactive compounds in Ajwa date fruit. Journal of agricultural and food chemistry. 2013;61(24):5834–40.

33. Sohilait MR, Pranowo HD, Haryadi W. Molecular docking analysis of curcumin analogues with COX-2. Bioinformation. 2017;13(11):356–9. doi: 10.6026/97320630013356. PubMed PMID: WOS:000449650400001.

34. Vo NNQ, Nomura Y, Muranaka T, Fukushima EO. Structure-Activity Relationships of Pentacyclic Triterpenoids as Inhibitors of Cyclooxygenase and Lipoxygenase Enzymes. J Nat Prod. 2019;82(12):3311–20. Epub 2019/11/28. doi: 10.1021/acs.jnatprod.9b00538. PubMed PMID: 31774676.

35. Xu S, Uddin MJ, Banerjee S, Duggan K, Musee J, Kiefer JR, et al. Fluorescent indomethacin-dansyl conjugates utilize the membrane-binding domain of cyclooxygenase-2 to block the opening to the active site. Journal of Biological Chemistry. 2019;294(22):8690–8. doi: https://doi.org/10.1074/jbc.RA119.007405.

36. Loza-Mejía MA, Salazar JR. Sterols and triterpenoids as potential anti-inflammatories: Molecular docking studies for binding to some enzymes involved in inflammatory pathways. Journal of Molecular Graphics and Modelling. 2015;62:18–25. doi: https://doi.org/10.1016/j.jmgm.2015.08.010.

